# A proteomics sample metadata representation for multiomics integration, and big data analysis

**DOI:** 10.1101/2021.05.21.445143

**Authors:** Chengxin Dai, Anja Füllgrabe, Julianus Pfeuffer, Elizaveta Solovyeva, Jingwen Deng, Pablo Moreno, Selvakumar Kamatchinathan, Deepti Jaiswal Kundu, Nancy George, Silvie Fexova, Björn Grüning, Melanie Christine Föll, Johannes Griss, Marc Vaudel, Enrique Audain, Marie Locard-Paulet, Michael Turewicz, Martin Eisenacher, Julian Uszkoreit, Tim Van Den Bossche, Veit Schwämmle, Henry Webel, Stefan Schulze, David Bouyssié, Savita Jayaram, Vinay Kumar Duggineni, Patroklos Samaras, Mathias Wilhelm, Meena Choi, Mingxun Wang, Oliver Kohlbacher, Alvis Brazma, Irene Papatheodorou, Nuno Bandeira, Eric W. Deutsch, Juan Antonio Vizcaíno, Mingze Bai, Timo Sachsenberg, Lev Levitsky, Yasset Perez-Riverol

**Affiliations:** Chongqing Key Laboratory on Big Data for Bio Intelligence, Chongqing University of Posts and telecommunications, Chongqing, China; European Molecular Biology Laboratory, European Bioinformatics Institute, Wellcome Genome Campus, Hinxton, UK; Algorithmic Bioinformatics, Freie Universität Berlin, Berlin, Germany; Visualization and Data analysis, Zuse Institute Berlin, Berlin, Germany; Moscow Institute of Physics and Technology, Dolgoprudny, Moscow Region, Russia; V.L. Talrose Institute for Energy Problems of Chemical Physics, N.N. Semenov Federal Research Center for Chemical Physics, Russian Academy of Sciences, Moscow, Russia; Bioinformatics Group Department of Computer Science, Albert-Ludwigs-University Freiburg, Freiburg, Germany; Institute for Surgical Pathology, Medical Center – University of Freiburg, Faculty of Medicine, University of Freiburg, Germany; Khoury College of Computer Sciences, Northeastern University, Boston, MA, USA; Department of Dermatology, Medical University of Vienna, Austria; Department of Clinical Sciences, University of Bergen, Norway; Department of Congenital Heart Disease and Pediatric Cardiology, Universitätsklinikum Schleswig-Holstein Kiel, Kiel, Germany; Novo Nordisk Foundation Center for Protein Research, University of Copenhagen, Denmark; Ruhr University Bochum, Medical Faculty, Medizinisches Proteom-Center, 44801 Bochum, German; Ruhr University Bochum, Center for Protein Diagnostics (PRODI), Medical Proteome Analysis, 44801 Bochum, German; VIB – UGent Center for Medical Biotechnology, VIB, Ghent, Belgium; Department of Biomolecular Medicine, Faculty of Medicine and Health Sciences, Ghent University, Ghent, Belgium; Department of Biochemistry and Molecular Biology, University of Southern Denmark, Campusvej 55, 5230 Odense, Denmark; University of Pennsylvania, Department of Biology, Philadelphia, PA 19104, USA; Institute of Pharmacology and Structural Biology, University of Toulouse, CNRS, UPS, France; nference Labs, Bengaluru, KA, 560017, India; Chair of Proteomics and Bioanalytics, Technical University of Munich, Germany; Skaggs School of Pharmacy and Pharmaceutical Sciences, University of California San Diego, La Jolla, CA 92093, USA; Biomolecular Interactions, Max Planck Institute for Developmental Biology, 72076, Tübingen, Germany; Institute for Bioinformatics and Medical Informatics, University of Tübingen, Sand 14, 72076, Tübingen, Germany; Department of Computer Science, University of Tübingen, WSI/ZBIT, Sand 14, 72076, Tübingen, Germany; Quantitative Biology Center, University of Tübingen, Auf der Morgenstelle 8, 72076, Tübingen, Germany; Institute for Translational Bioinformatics, University Hospital Tübingen, Sand 14, 72076, Tübingen, Germany; Center for Computational Mass Spectrometry, Department of Computer Science and Engineering, Skaggs School of Pharmacy and Pharmaceutical Sciences, University of California, San Diego, 92093-0404, USA; Institute for Systems Biology, 401 Terry Ave N, Seattle, WA, 98109, USA; State Key Laboratory of Proteomics, Beijing Proteome Research Center, National Center for Protein Sciences (Beijing), Beijing Institute of Life Omics, Beijing 102206, China

## Abstract

The amount of public proteomics data is increasing at an extraordinary rate. Hundreds of datasets are submitted each month to ProteomeXchange repositories, representing many types of proteomics studies, focusing on different aspects such as quantitative experiments, post-translational modifications, protein-protein interactions, or subcellular localization, among many others. For every proteomics dataset, two levels of data are captured: the dataset description, and the data files (encoded in different file formats). Whereas the dataset description and data file formats are supported by all ProteomeXchange partner repositories, there is no standardized format to properly describe the sample metadata and their relationship with the dataset files in a way that fully allows their understanding or re-analysis. It is left to the user’s choice whether to provide or not an *ad hoc* document containing this information. Therefore, in many cases, understanding the study design and data requires going back to the associated publication. This can be tedious and may be restricted in the case of non-open access publications. In many cases, this problem limits the generalization and reuse of public proteomics data.

Here we present a standard representation for sample metadata tailored to proteomics datasets produced by the HUPO Proteomics Standards Initiative and supported by ProteomeXchange resources. We repurposed the existing data format MAGE-TAB used routinely in the transcriptomics field to represent and annotate proteomics datasets. MAGETAB-Proteomics defines a set of annotation rules that the datasets submitted to ProteomeXchange should follow, ranging from sample properties to data analysis protocols. We also introduce a crowdsourcing project that enabled the manual curation of over 200 public datasets using MAGE-TAB-Proteomics. In addition, we describe an ecosystem of tools and libraries that were developed to validate and submit sample metadata-related information to ProteomeXchange. We expect that these tools will improve the reproducibility of published results and facilitate the reanalysis and integration of public proteomics datasets.

## Introduction

The amount of proteomics data in public repositories is growing at an unprecedented rate ^1, 2^. ProteomeXchange (PX) is a consortium of multiple proteomics resources including the PRIDE database ^2^, PASSEL and PeptideAtlas ^3^, MassIVE ^4^, jPOST ^5, 6^, iProX ^7^, and Panorama Public ^8^. As of May 2021, over 25,000 datasets have been submitted to ProteomeXchange data repositories. Ranging from peptide and protein identification to proteome turnover experiments, ProteomeXchange datasets cover the whole spectrum of mass spectrometry (MS) analytical methods and experimental designs, which enable biologists and clinicians to study different aspects of the proteome. In parallel to the generalisation of data deposition, data reuse of public datasets has become increasingly very popular for different purposes. However, thus far been, it has been mainly limited to benchmarking studies and/or applications related to peptide and protein identification, with resources such as PeptideAtlas ^3^ and GPMDB ^9^ systematically reanalyzing data from ProteomeXchange^10^. Recently, new efforts like ProteomicsDB ^11^, MassiVE.Quant ^4^ and Expression Atlas ^12^ have started to include reanalysed quantitative public datasets to present baseline and differential protein expression. However, the scalability and broad reuse of public quantitative experiments have been limited by the lack of sample metadata annotation, which unambiguously associates the samples included in each dataset with the corresponding data files ^13, 14^.

Since 2012 ProteomeXchange resources have been capturing a general dataset description including dataset title, description, instrument, protein modifications included in the search, and submitters/principal investigators ^1^. The files included in each dataset are, on one hand, the output of the corresponding instrument (e.g., raw files), and on the other, the processed results, which can be represented e.g., in standard file formats such as mzIdentML ^15^ or mzTab ^16^. Currently, all ProteomeXchange partners mandate two types of information for each dataset: a general dataset description and the files containing the different required data types. Unfortunately, the experimental design and sample-related information are often missing in the datasets or are stored in *ad hoc* ways and formats ^1^. Information about the biological samples such as organism parts, disease, or cell lines, and the links between the samples and the corresponding data files are mostly missing.

Sample-related metadata and their relationship with the data files are well-captured in two widespread file formats called ISA-TAB ^17^ and MAGE-TAB ^18^, which are used in other omics fields such as metabolomics and transcriptomics, respectively. As of May 2021, ArrayExpress has stored over 74,000 functional genomics datasets in the MAGE-TAB format ^18, 19^. In both formats, a tab-delimited file is used to annotate the sample metadata and link the metadata to the corresponding data file(s). In MAGE-TAB this information is encoded in two different files: the Sample and Data Relationship File (SDRF) format and the Investigation Description File (IDF) format, which stores the general dataset information. While MAGE-TAB was originally designed for microarray experiments, it has been successfully adapted to high-throughput RNA-sequencing and single-cell RNA-Seq experiments^20^.

Here we introduce an extension and implementation of the MAGE-TAB format for proteomics (MAGE-TAB-Proteomics). The format has been developed in collaboration with the Proteomics Standards Initiative (PSI), the organisation in charge of developing open standard formats in the field ^21^. We have also developed general guidelines about what information needs to be encoded in the SDRF part of the MAGE-TAB to improve the reproducibility and enable the reanalysis of proteomics datasets. These include some information that is already required to submit a dataset to ProteomeXchange, including sample attributes (e.g., organism part), but also analytical information such as fraction-related information and labelling channels (e.g., TMT, SILAC), which are required to understand the experimental design. In addition, we have crowdsourced the annotation of over 200 existing public datasets according to these guidelines. These cover different analytical methods and experimental designs, ranging from label-free quantification to affinity purification mass spectrometry experiments (AP-MS). Finally, we have developed an ecosystem of tools to validate MAGE-TAB-Proteomics files and integrate the metadata in the PRIDE database, the most popular ProteomeXchange resource. The full specification document describing in detail all aspects of version 1.0 of the MAGE-TAB-Proteomics standard is available at the PSI website (http://psidev.info/magetab) together with a detailed description of the current implementations, including examples.

## MAGE-TAB for Proteomics

### IDF: Providing study description information

The IDF file contains information describing the study, including e.g., authors/submitters, protocols, and publications (**Supplementary Note 1**). The IDF format contains a series of key/value pairs, where each key represents a different property. For example, “Experiment Description” should be followed by a free-text description of the experiment (which would be the value). Most of the fields can contain more than one value, so that multiple values (for example, multiple analysis software tools) can be defined in a single IDF file. Since 2012 ProteomeXchange resources have used an XML file format for describing datasets, called PX XML (http://proteomecentral.proteomexchange.org/schemas/proteomeXchange-1.4.0.html), that captures equivalent information to the ones included in IDF, making both files easily exchangeable (**Supplementary Note 1**). Therefore, we simply defined MAGE-TAB IDF based on the existing PX XML format. Here we will mainly focus on the formalization and standardization of the new MAGE-TAB-Proteomics SDRF.

### SDRF: Linking samples to data files

SDRF is a tab-delimited file that describes the samples and allows their mapping to the data files ^1^. As shown in Figure 1a, SDRF includes the annotation of: (i) biological sample metadata; (ii) the relationships between samples and data files; (iii) the raw data files’ metadata; and (iv) the variables under study (called factor values). Each row in an SDRF file corresponds to one relationship between a sample and a data file (an MS raw file, or a channel included in a given raw file in the case of labelling). Each column corresponds to an attribute/property of the sample or the file, and the value in each cell is the specific value of the property (**Figure 1a**).

**Figure 1:**
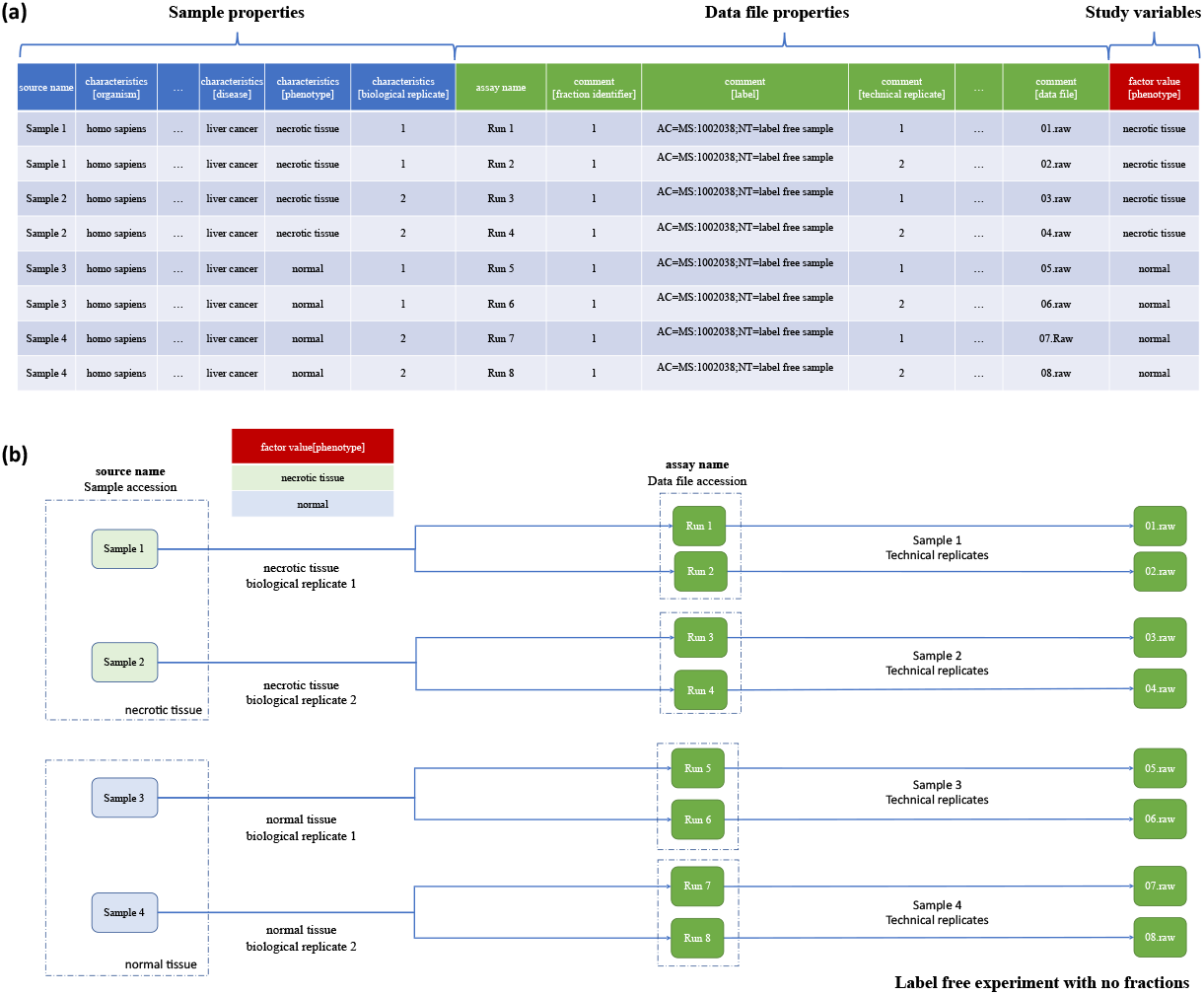
SDRF-Proteomics representation for a label-free-based experiment. **(a)** the SDRF tab-delimited file including the three main sections highlighted: sample metadata, data file properties, and the variables under study (factor values). **(b)** a label-free experimental design including two biological replicates and two technical replicates per biological replicate. The biological and technical replicates are defined by the variable under study (e.g., phenotype).

All the properties in the SDRF must be encoded as ontology terms whereas the values of the properties can be encoded as ontology terms, numerical values, or free text. To facilitate the annotation, validation, and processing of SDRF files, we have defined in the specification document a list of ontologies that can be used for encoding each property. For example, most of the sample properties are included in the Experimental Factor Ontology ^22^ (EFO - https://www.ebi.ac.uk/efo/), while most of the data file properties are included in PSI-MS controlled vocabulary (https://www.ebi.ac.uk/ols/ontologies/ms) and the PRIDE ontology (https://www.ebi.ac.uk/ols/ontologies/pride).

Each sample in an SDRF file has a unique identifier (source name), and every sample property is encoded using the prefix *characteristics* (e.g., *characteristics [organism part]*). Conversely, each data file has a unique identifier (assay name), and every file property has the prefix *comment* (e.g., *comment[instrument], comment[fraction identifier]*). Finally, the variables under study must be specified with the prefix *factor value* (*e.g., factor value[tissue]*). The MAGE-TAB-Proteomics specification defines the minimum information that should be provided for every sample and data file (see specification document). For all proteomics experiments, the following properties must be provided: organism, organism part, and biological replicate accession. For every data file, the following properties are required: fraction identifier, technical replicate accession, label (labelling method), and data file name. Biological and technical replicates should be explicitly included using the terms *characteristics[biological replicate]* and *comment[technical replicate]*, respectively (**Figure 1a**). The biological replicate field is considered a property of the samples, whereas the technical replicate is considered a property of the data files.

A second category of fields includes pieces of information that are not mandatory but recommended. This comprises properties of the data files such as instrument model, cleavage agents, or mass modifications, which include post-translational modifications (PTMs) of biological interest, artefactual mass modifications considered in the analysis, and labelling. Most of the values of these properties can be encoded as a combination of multiple key/value pairs (e.g. to specify methionine oxidation: AC=UNIMOD:35;NT=Oxidation;MT=Variable;TA=M).

Each repository can define which of the recommended fields must be provided in their resource, depending on the experiment types. We have defined the concept of templates (**Supplementary Note 2**), which form a set of recommended properties required per experiment type. Submitters can begin using the template corresponding to their experiment, to streamline annotation. For example, for cell line experiments, the cell line (*characteristics[cell line]*) is a recommended metadata item. Another example is that for every human dataset, the disease under study should be provided in the field *characteristics[disease]* and the control samples should be labelled with the value “normal”. The ProteomeXchange templates request for the submitters to provide for every data file the following properties: instrument model, cleavage agent, mass modifications, fragment mass tolerance, and precursor mass tolerance. We believe that this is the minimum set of information that is necessary for understanding and properly analysing MS-based proteomics datasets.

### Multiplexing and fractionation

In contrast to transcriptomics datasets, where a one-to-one relationship between each sample and data file is the predominant pattern, there are two experimental designs in proteomics that do not follow this trend: sample multiplexing and fractionation (**Figure 2a-b**). As mentioned above, in quantitative experiments involving labelling and multiplexing (e.g., tandem mass tag (TMT) datasets), multiple samples can be related to the same data file (**Figure 2a**). In those cases, the data file properties should be repeated for each sample including all the properties (e.g., instrument), and the relevant property labels (e.g., comment[label] = TMT128N) can be used to encode the different samples included in the same data file. In contrast, where fractionation is used, the sample information should be repeated for each data file. A property called fraction identifier is used to make clear which fractions correspond to a given data file (e.g., *comment[fraction identifier]* = 1). As a result, MAGE-TAB-Proteomics is highly flexible and can be applied to complex experimental designs such as the one presented in **Figure 2b**. This figure depicts a TMT experimental design, where three samples are fractionated three times each. The resulting SDRF contains nine rows, where samples are repeated for each of their fractions, and the data file properties are repeated for each sample channel, which is encoded using the property *comment[label]*.

**Figure 2:**
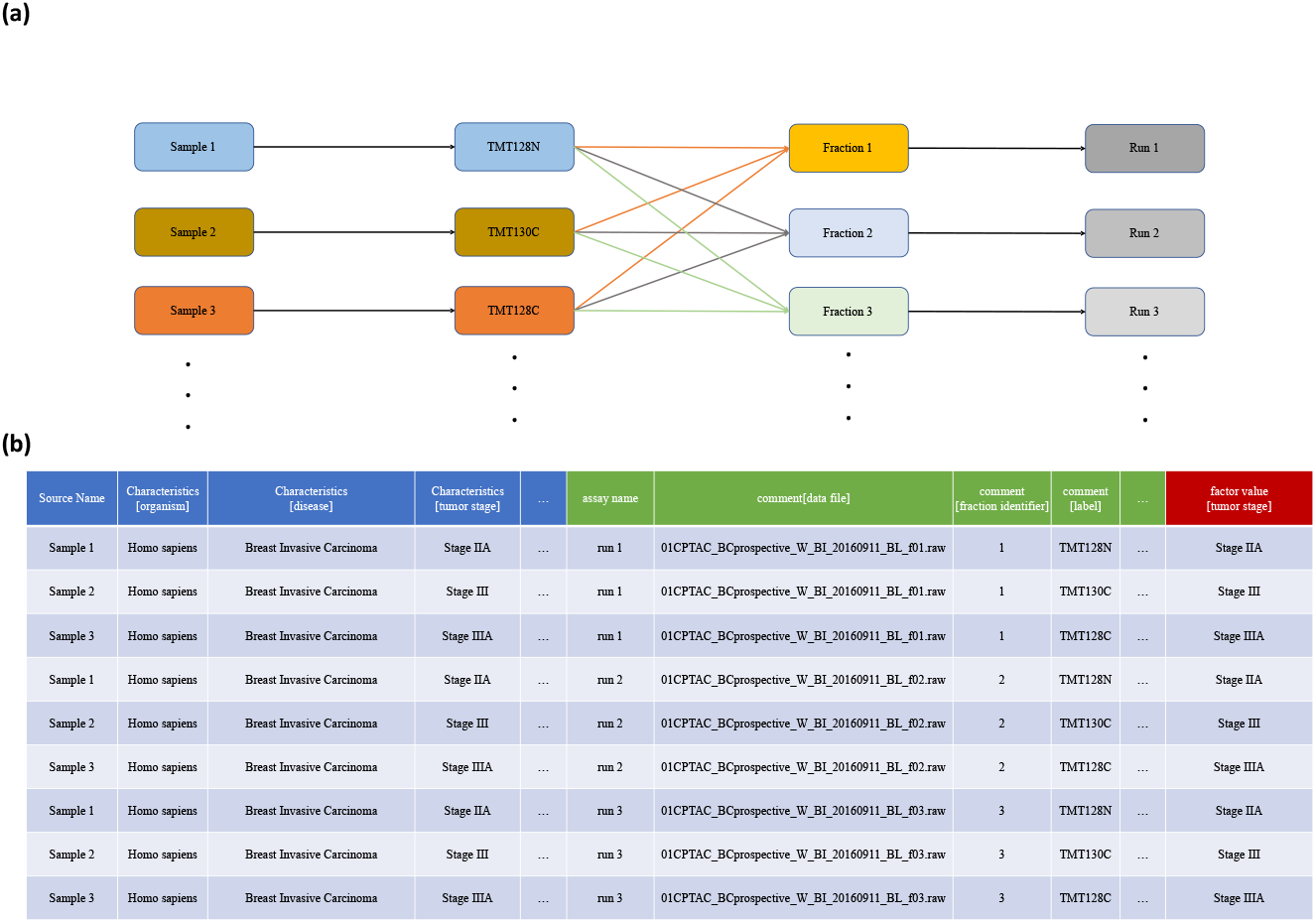
SDRF-Proteomics file corresponding to a TMT experiment. (**a**) TMT experiment design with three samples and three fractions. (**b**) SDRF representation for a TMT experiment with three samples and 3 fractions resulting in nine rows where samples are repeated for each fraction and data the file information is repeated for each labelling channel.

While the duplication of information can be perceived as redundant, it enables a streamlined data re-analysis because each row/line of the SDRF can be processed individually. In addition, it facilitates meta-analysis with simple operations such as merging different SDRFs coming from different datasets or splitting an SDRF for a given dataset by a specific property of the data file or the sample.

### Crowdsourcing annotations of public proteomics datasets

Crowdsourcing has been successfully applied to address key problems in bioinformatics, yielding a series of success stories ^23^. The European Bioinformatics Community for Mass Spectrometry (EuBIC-MS ^24^, https://eubic-ms.org) set up a collective effort to annotate existing ProteomeXchange datasets using MAGE-TAB-Proteomics ^14^. Volunteers from multiple institutes joined the task of annotating, discussing, and improving the SDRF format, openly and collaboratively. Each annotator could create an issue in GitHub about a project of interest, fork the main project repository (https://github.com/bigbio/proteomics-metadata-standard), and annotate the corresponding dataset locally on their computers. Then, a pull request was submitted to include the added annotations. An independent group of reviewers checked that the proposed SDRF conformed to the guidelines before being approved. This collaborative peer review system allowed the identification of potential issues in the file format, which is now compatible with the main MS experiment types. Additionally, a file validator was developed (https://github.com/bigbio/sdrf-pipelines) to automatically perform a semantic and structural validation of new SDRF files. Table 1 shows a group of gold standard annotated datasets from ProteomeXchange that can be utilized as examples when creating an SDRF file for label-free experiments, multiplex TMT and SILAC datasets as well as AP-MS and phospho-enriched samples.

**Table 1:** Examples of MAGE-TAB-Proteomics files.

| Dataset Type | ProteomeXchange / PubMed Accession | MAGE-TAB |
| --- | --- | --- |
| Label-free | PXD008934 | https://github.com/bigbio/proteomics-metadata-standard/tree/master/annotated-projects/PXD008934 |
| TMT, CPTAC dataset not in ProteomeXchange | CPTAC<br>PMID27251275 | https://github.com/bigbio/proteomics-metadata-standard/tree/master/annotated-projects/PMID33212010 |
| SILAC | PXD006877 | https://github.com/bigbio/proteomics-metadata-standard/tree/master/annotated-projects/PXD006877 |
| Phospho-proteomics | PXD006482 | https://github.com/bigbio/proteomics-metadata-standard/tree/master/annotated-projects/PXD006482 |
| Label-free, multiple fragmentation modes and various enzymes | PXD010154 | https://github.com/bigbio/proteomics-metadata-standard/tree/master/annotated-projects/PXD010154 |
| AP-MS interactomics | PXD018117 | https://github.com/bigbio/proteomics-metadata-standard/tree/master/annotated-projects/PXD018117 |
| TMT | PXD017710 | https://github.com/bigbio/proteomics-metadata-standard/tree/master/annotated-projects/PXD017710 |
| Label-free | PXD004242 | https://github.com/bigbio/proteomics-metadata-standard/tree/master/annotated-projects/PXD004242 |

As of May 2021, over 200 public datasets have been annotated, covering a broad spectrum of organisms, quantitative and data acquisition methods, as well as enrichment/fractionation strategies (**Figure 3).** However, for most multiplexed experiments available on ProteomeXchange, the sample to label assignment was not specified by the authors since it was not possible to map which channel was associated with which sample in a standardized format. As a consequence, such datasets are underrepresented in the collection of annotated datasets. The lack of such essential experimental design information precludes the reprocessing and reproduction of quantitative results. This highlights the urgent need for systematic and standardized metadata annotations.

**Figure 3:**
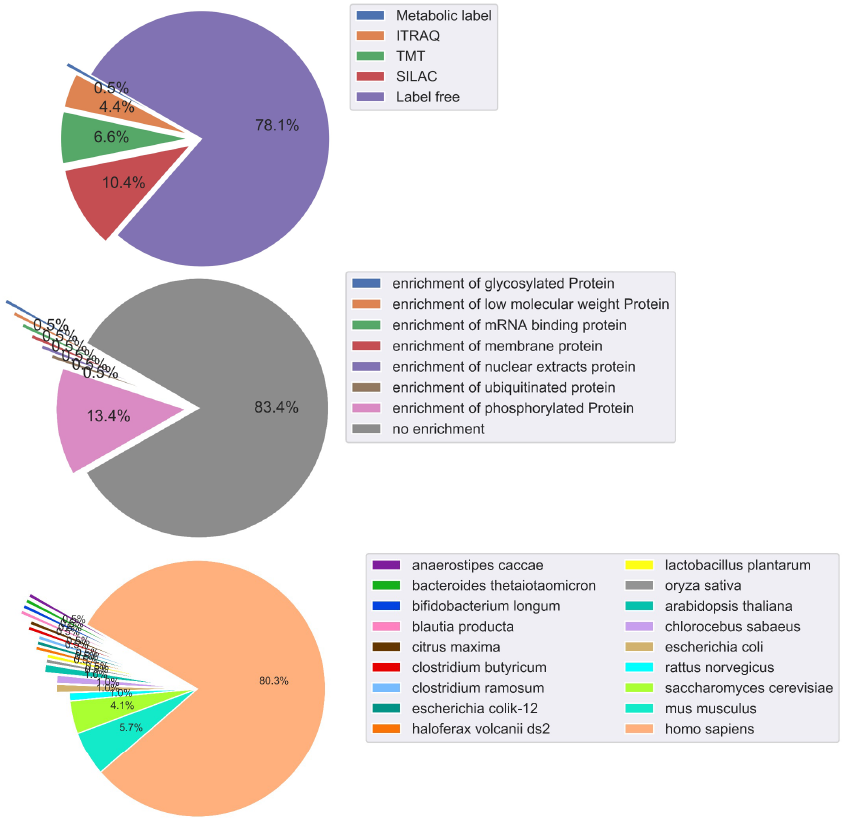
The distribution of the 200 annotated projects in organisms, quantitative methods, data acquisition methods, and enrichment methods. Projects are classified into three different categories: analytical method including Label-free, TMT, SILAC, or ITRAQ experiments; enrichment process including phosphorylation and glycosylation; and project species.

### The MAGE-TAB-Proteomics toolbox

In order to facilitate automatic data analysis and reuse, we developed a set of software libraries that enable the validation and conversion from and to the MAGE-TAB-Proteomics format to parameter files (**Figure 4a**). For transcriptomics data, the main library to validate a MAGE-TAB file is the Bio-MAGETAB package (https://metacpan.org/release/Bio-MAGETAB). We implemented a Java and a Python library to enable the validation of MAGE-TAB-Proteomics files, and in particular of the SDRF-Proteomics part.

**Figure 4:**
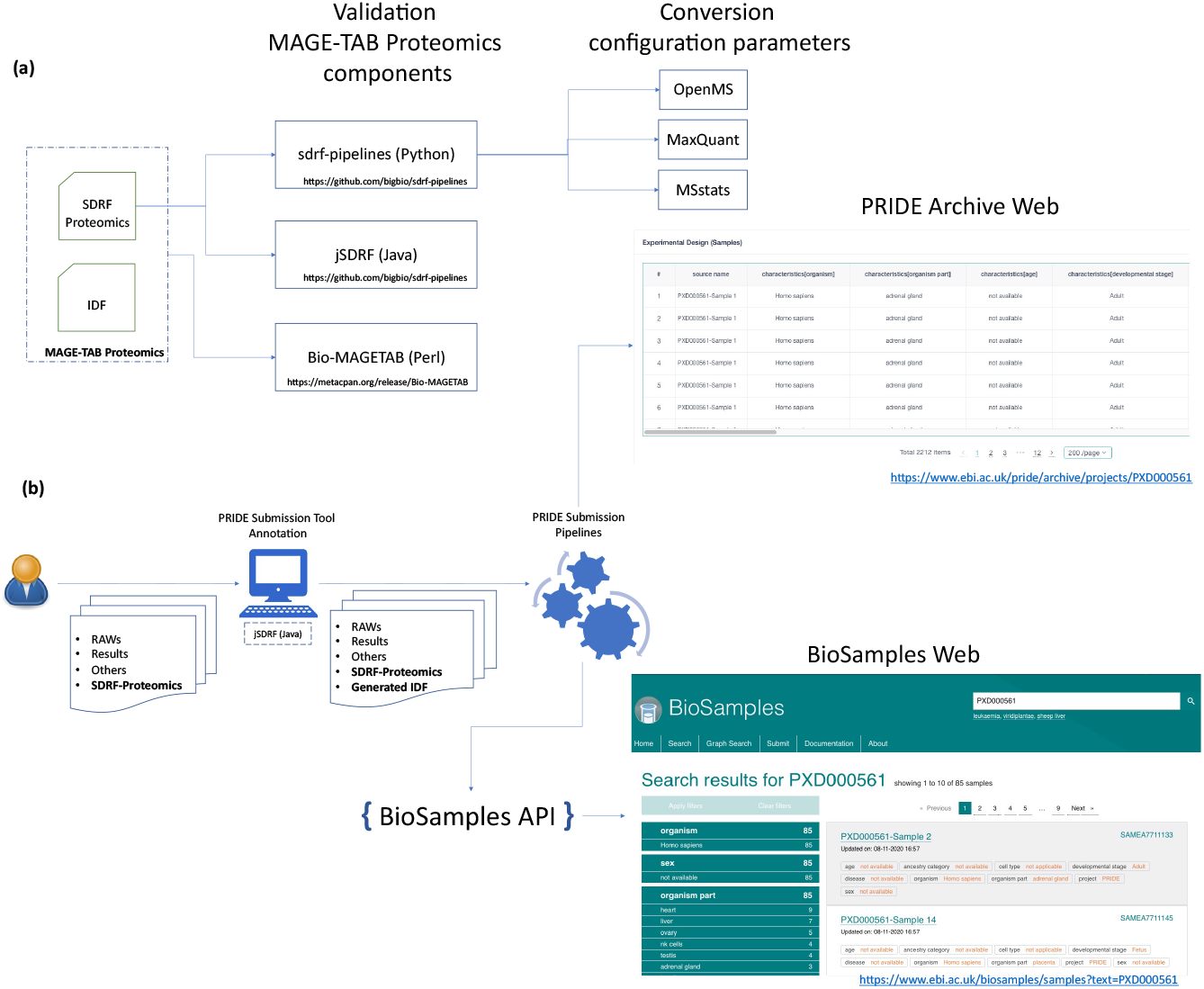
(**a**) MAGE-TAB for proteomics toolbox and libraries in different languages including Python, Java, and Perl. The libraries provide two main functionalities validation and conversion of the sample metadata SDRF different configuration files of popular proteomics tools like MaxQuant, MSstats, or OpenMS. (**b**) PRIDE database submission workflow supports SDRF proteomics files. The file format can be provided in the Submission tool, validated by the submission pipelines, and finally, the sample information is shown on the Web page of each dataset. The sample information is submitted to the BioSample database at EBI and a unique accession is assigned by that resource to each sample.

A Python package called sdrf-pipelines (https://github.com/bigbio/sdrf-pipelines) enables the validation of the structure and also of the semantic rules applied to SDRF. It validates the files according to different experiments and data types, defined in the templates.

For example, if the template corresponds to a human dataset, the software validates that the sample metadata complies with the human template: organism, disease, ancestry, etc. The sdrf-pipelines package can be installed from different package managers like BioConda ^25^, or BioContainers ^26^. In addition, it allows users to convert SDRF files to other popular proteomics software’s configuration/input files such as MaxQuant ^27^, OpenMS ^28^, and MSstats ^29^, to facilitate the automation of dataset reanalyses. The counterpart Java library, called jsdrf (https://github.com/bigbio/isdrf), similarly enables the validation of SDRF files. It also includes a generic data model that can be used by Java applications to validate and handle SDRF-Proteomics files.

### Submission of proteomics datasets to the PRIDE database

In PRIDE, datasets are submitted using the ProteomeXchange submission tool, a desktop application that guides the users through a set of steps to construct the submission and finally performs the transfer of the data files^2^. The submission tool enables annotation of the submitted project, including, among others, the title of the dataset, a general description, and sample and data analysis protocols (**Figure 4**). In the final step, all the files associated with the dataset are added to the submission, including the MS raw files, peptide and protein identification results files, and optionally, other additional file types. All the information annotated by the submitter is encoded into a *submission.px* file that is now converted to the MAGE-TAB-Proteomics IDF automatically using PRIDE internal pipelines once the submission is received and processed.

Since May 2021, the submission tool accepts SDRF as a supported file. These can be created externally and submitted with the dataset. It is then recognized as an ‘EXPERIMENT DESIGN’ type (**Figure 4**). As mentioned above, the library jSDRF is used to validate the files. For example, the library can validate if the SDRF contains each sample and data file relationship, the labelling, fraction identifiers, and sample and data accessions. PRIDE internal pipelines can then be triggered to create MAGE-TAB-Proteomics files containing the IDF (converted from *submission.px*) and the SDRF provided by the user. Then, the metadata assigned to each sample is submitted to the BioSamples database^30^ and a BioSample accession number is created for each sample in the experiment (**Figure 4b**), which enables the linking between samples included in multi-omics datasets. The PRIDE web interface then presents for each dataset the associated sample metadata table (**Figure 4b**). In addition, the sample metadata is indexed by PRIDE, allowing users to locate and link samples and experiments within the vast number of public datasets.

### Conclusions and future perspectives

Recently, multiple resources have started to systematically reanalyze proteomics public quantitative datasets, including MassIVE.Quant, ExpressionAtlas and ProteomicsDB ^4, 11, 19^. However, the lack of sample metadata that enables the association between samples and data files and the corresponding properties of the sample, makes this task complex and unscalable. MAGE-TAB-Proteomics builds upon a popular and flexible data format in transcriptomics to represent proteomics experimental designs. Ranging from peptide/protein identification, analytical methods, affinity purification MS experiments, to differential abundance approaches, MAGE-TAB Proteomics enables the annotation of datasets generated using different experimental approaches. MAGE-TAB-Proteomics has a great potential to facilitate the automatic re-analysis of all currently available and future annotated public proteomics datasets with standardized workflows, and thereby enormously increase their inherent value. For such large-scale re-analyses, the use of relatively large hardware resources in cloud environments will be necessary.

The proposed standard representation will facilitate the integration and annotation of proteomics studies, thereby enhancing their discoverability, reproducibility, and amenability to systematic reprocessing. In addition, it will enable the development of common submission systems of multi-omics datasets between resources different types of omics data, or even new multi-omics resources, enabling submission, validation, and visualization of multi-omics data^31, 32^. Multiple efforts are ongoing to provide tools that help users to annotate datasets on the web. Novel use cases and more types of MS-based experiments are being discussed by the MAGE-TAB-Proteomics team. At the moment of writing, the representation in MAGE-TAB-Proteomics of more experimental approaches is actively under discussion, including MS imaging and xenograft proteomics experimental designs. This number is expected to keep growing in the future, as experts in additional proteomics experimental approaches can be included in the discussion. This is an analogous process to what has happened with other PSI formats in the past. Additionally, the format has also already triggered some interest in the MS metabolomics field. By May 2021, multiple metabolomics datasets coming from the ReDU resource ^33^ have been added to the annotation repository.

Importantly, due to the gradual and iterative implementation of the standard into ProteomeXchange submission pipelines and related tools, the adoption of the standard will not create a major additional burden for users. We expect that this initiative will therefore improve reproducibility, facilitate the development of new tools dedicated to proteomics data analysis, and facilitate data re-analysis efforts and related resources, making the re-use of proteomics data more popular for biologists (analogously to what has happened in e.g., the transcriptomics field). It is also key that this effort is supported jointly by the world-leading proteomics databases in ProteomeXchange. Through our open, collaborative, and community-based approach, we invite all parties interested to join the initiative.

## Supporting information

HUPO PSI Specification

## Acknowledgements

YPR, SK, DJK, and JAV would like to acknowledge funding from the Wellcome Trust grant number 208391/Z/17/Z and EMBL core funding. MLP is supported financially by the Novo Nordisk Foundation (Grant agreement NNF14CC0001). MT is supported by de.NBI, a project of the German Federal Ministry of Education and Research (BMBF) [grant number FKZ 031 A 534A]. ME and JU are members of the Center for Protein Diagnostics (PRODI), a grant from the Ministry of Innovation, Science, and Research of North-Rhine Westphalia, Germany. TVDB is supported by the Research Foundation – Flanders (SB grant 1S90918N). EWD acknowledges NIGMS grants R01GM087221, R24GM127667, and NSF grants 1933311 and 1922871. Funds for the overall project were also made available by an ELIXIR Implementation Study. CD, MB are supported by the National Key Research and Development Program of China (2017YFC0908404, 2017YFC0908405) and the Natural Science Foundation of Chongqing, China (cstc2018jcyjAX0225).

## Contributions

Y.P.R, C.D, J.D, E.S, D.J.K, M.C.F, J.G, M.V, E.A, M.L.P, M.T, M.E, J.U, T.V.D.B, V.S, H.W, S.S, D.B, S.J, V.K.D, P.S, M.W, A.F, N.G, S.F, M.C, M.B, O.K, A.B, I.P, N.B, E.W.D, J.A.V contributed to the definition of the specification and annotated the GitHub public proteomics datasets. Y.P.R, C.D, J.P, P.M, S.K, B.G, L.L and T.S developed the MAGE-TAB for proteomics libraries and integration with PRIDE database. N.B, M.C, E.W.D, J.A.V, Y.P.R introduced the specification to HUPO-PSI and ProteomeXchange consortium. All authors contributed with the writing and review of the present manuscript.

## Abbreviations

AP-MS: Affinity Purification coupled to Mass Spectrometry
EuBIC-MS: European Bioinformatics Community for Mass Spectrometry
IDF: Investigation Description File
PSI: Proteomics Standards Initiative
MAGE-TAB: MicroArray Gene Expression Tabular
MS: Mass Spectrometry
PRIDE: PRoteomics IDEntification Database
PX: ProteomeXchange
SDRF: Sample and Data Relationship Format
TMT: Tandem Mass Tag

