## Supplementary Information for "A proteomics sample metadata representation for multiomics integration, and big data analysis"

### **Supplementary Note 1: Investigation description format**

The IDF (Investigation Description Format) file contains fields describing the study, authors/submitters, protocols, publications (Read Section). ProteomerXchange resources developed a file format called *submission.px* which captures the same information as the MAGE-TAB IDF. We have developed a simple Python tool that converts each submission in PRIDE Archive database to IDF (<https://github.com/bigbio/proteomics-metadata-standard/blob/master/generate_idf.py>).

Figure 1 (<https://emblebi-my.sharepoint.com/:p:/g/personal/yperez_ebi_ac_uk/EcLv3kR8-4BFp_qnK43eM2UB27e4wWNARY6xRbVKVNMXBQ?e=wfcBI3>): IDF for ProteomeXchange project PXD000612.

Figure 1 shows an example of a submission.px and their corresponding IDF. The IDF component of a MAGE-TAB document consists of a set of unique tags attached to their corresponding values in a simple tab-delimited text format. For example, "Experiment Description" should be followed by a free-text description of the experiment. Most of the following fields can be used with more than one value, so that (for example - multiple experimental factors etc) multiple values can be defined in a single IDF file. In such cases the multiple values should be separated by semicolons (";")

Some of the tags of the IDF are:

**Investigation Title**: The overall title of the investigation. This tag can only have one value.

Corresponding field in proteomeXchange.xml: Title

**Experiment Description:** A short paragraph describing the experiment as free-text. This tag can only have one value. The text should clearly explain what you did in your experiment - this will help the curation team to check and process your MAGE-TAB document.

Corresponding field in proteomeXchange.xml: Description

**Date of Experiment**: The date on which the experiment was performed. This tag can only have one value. Some databases like PRIDE provides the Submission data which can be consider as the Date of the Experiment.

**Public Release Date:** The date on which the experimental data will be/was released. You can ask us to change this later. This tag can only have one value.

Corresponding field in proteomeXchange.xml: announceDate

**Person details**: The proteomics community captures Person details differently than the IDF MAGE-TAB specification. The Person information is captured in ProteomeXchange as a list of contacts where each contact is a list of CvTerms. The name of the CVterm is the name of the attribute, value of the CvTerm is the value of the attribute. For example, in <cvParam cvRef="MS" accession="MS:1000586" name="contact name" value="Christoph Krisp"/> the name of the CVTerm is the contact name, the value is the name of the person.

In the IDF, a Contact is a Person with different properties, for example:

- Person Last Name: The last name of each person associated with the experiment.
- Person First Name: The first name of each person associated with the experiment.
- Person Email: The email address of each person associated with the experiment.
- Person Affiliation: The organization affiliation for each person associated with the experiment. This tag is mandatory for sequencing submissions.
- Person Roles: The role(s) performed by each person. Typically, these terms should come from the Experimental Factor Ontology. See for example the list of organization role terms. If more than one role is needed per person, the roles should be given as a semicolon (;) delimited list. The roles defined by ProteomeXchange are two: dataset submitter; or lab head
- Person Roles Term Source REF: The source of the Person Roles terms; his must reference one of the Term Source Names defined in the IDF file.
- Person Roles Term Accession Number: The accession number for this term, taken from the indicated Term Source.

**Publication Details**:

- PubMed ID: The PubMed IDs of the publication(s) associated with this investigation (where available)
- Publication DOI: A Digital Object Identifier (DOI) for each publication (where available).
- Publication Author List: The list of authors associated with each publication.
- Publication Title: The title of each publication.
- Publication Status: A term describing the status of each publication (e.g. submitted, in preparation, published).
- Publication Status Term Source REF: The source of the Publication Status terms; his must reference one of the Term Source Names defined in the IDF file.
- Publication Status Term Accession Number: The accession number for this term, taken from the indicated Term Source.

**Sample and Data Protocols**: The sample and data protocols in transciptomics are captured with a low-level details, while in proteomics is a summary of multiple protocols within two categories: Sample and Data Protocols. For that reason, we recommend writing the sample and data protocols in the following standard

- Sample Protocol: Protocol Name, The names of the protocols used within the MAGE-TAB document. The sample protocol name for PX submissions will be: *P-MTAB-Sample-PXID*. The protocol name will be the combination of Sample and the Submission PX in ProteomeXchange
- Protocol Type: The type of the protocol, taken from a controlled vocabulary. Typically, this term should come from the Experimental Factor Ontology . See for example the list of protocol terms. The protocol type for PX submissions will be: sample collection protocol
- Protocol Description: A free-text description of the protocol. This text is included in a single tab-delimited field. The Protocol Description is the present Sample Description in ProteomeXchange.
- Protocol Parameters: A semicolon-delimited list of parameter names.
- Protocol Hardware: The protocol hardware is the instrument that was use to capture the sample. If multiple instruments are used, they should be separated by (;)
- Data Protocol: The Data protocol is a generic way in proteomics to capture all the metadata about the data analysis steps.
- Protocol Software: The software used by the protocol.

**Experimental Factors**:

- Experimental Factor Name: A user-defined name for each experimental factor studied by the experiment. These experimental factors represent the variables within the investigation (e.g. growth condition, genotype, organism part). The actual values of these variables will be listed in the SDRF file, in "Factor Value [<factor name>]" columns.
- Experimental Factor Type: A term describing the type of each experimental factor. These terms will usually come from the Experimental Factor Ontology.
- Experimental Factor Term Source REF: The source of the Experimental Factor Type terms; this must reference one of the Term Source Names defined in the IDF file.
- Experimental Factor Term Accession Number: The accession number for this term, taken from the indicated Term Source.

**SDRF File**: The name(s) of the SDRF file(s) accompanying this IDF file.

**Additional Properties**:

- ProteomeXchange accession number: Main identifier of a ProteomeXchange dataset.

**Supplementary Note 2: Templates of SDRF for ProteomeXchange repositories**

**Sample attributes**: Minimum sample attributes for primary cells coming from different species and cell lines

|  | Default | Human | Vertebrates | Non-vertebrates | Plants | Cell lines | Ontology/CV URL |
| --- | --- | --- | --- | --- | --- | --- | --- |
| Source Name | 1 | 1 | 1 | 1 | 1 | 1 |  |
| characteristics[organism] | 1 | 1 | 1 | 1 | 1 | 1 | <https://www.ebi.ac.uk/ols/ontologies/ncbitaxon> |
| characteristics[strain/breed] | 0 | 0 | 0 | 0️ | 0 | 0️ | <https://www.ebi.ac.uk/ols/ontologies/efo> |
| characteristics[ecotype/cultivar] | 0 | 0 | 0 | 0 | 0️ | 0 | <https://www.ebi.ac.uk/ols/ontologies/efo> |
| characteristics[ancestry category] | 0 | 1 | 0 | 0 | 0 | 0 | <https://www.ebi.ac.uk/ols/ontologies/efo> |
| characteristics[age] | 0 | 1 | 0️ | 0 | 0️ | 0 |  |
| characteristics[sex] | 0 | 1 | 0️ | 0 | 0 | 0 | <https://www.ebi.ac.uk/ols/ontologies/efo> |
| characteristics[disease] | 1 | 1 | 1 | 1 | 0 | 1 | <https://www.ebi.ac.uk/ols/ontologies/efo> |
| characteristics[organism part] | 1 | 1 | 1 | 1 | 1 | 1 | <https://www.ebi.ac.uk/ols/ontologies/efo> |
| characteristics[cell type] | 1 | 1 | 1 | 1 | 1 | 1 | <https://www.ebi.ac.uk/ols/ontologies/efo> |
| characteristics[individual] | 0 | 1 | 0️ | 0️ | 0️ | 0️ |  |
| characteristics[cultured cell] | 0 | 0 | 0 | 0 | 0 | 1 | <https://www.ebi.ac.uk/ols/ontologies/efo> |
| characteristics[biological replicate] | 1 | 1 | 1 | 1 | 1 | 1 |  |
| comment[data file] | 1 | 1 | 1 | 1 | 1 | 1 |  |
| comment[fraction identifier] | 1 | 1 | 1 | 1 | 1 | 1 |  |
| comment[label] | 1 | 1 | 1 | 1 | 1 | 1 | <https://www.ebi.ac.uk/ols/ontologies/ms>  <https://www.ebi.ac.uk/ols/ontologies/pride> |
| comment[cleavage agent details] | 1 | 1 | 1 | 1 | 1 | 1 | <https://www.ebi.ac.uk/ols/ontologies/ms> |
| comment[instrument] | 1 | 1 | 1 | 1 | 1 | 1 | <https://www.ebi.ac.uk/ols/ontologies/ms> |
| comment[technical replicate] | 1 | 1 | 1 | 1 | 1 | 1 |  |

1: Required Attributes for each sample type (e.g., human, vertebrates).

0: Optional Attributes
