## Supplementary material for "A proteomics sample metadata representation for multiomics integration, and big data analysis": HUPO PSI Specification

PSI Recommendation

PSI Mass Spectrometry and Proteomics Informatics Working Groups

Status: DRAFT

Date: 25 March 2021

Title: MAGE-TAB Proteomics: Sample and Data Relationship Format (SDRF) for Proteomics

Chengxin Dai, Chongqing University of Posts and Telecommunications, Chongqing, China

Anja Fullgrabe, EMBL-EBI, U.K

Elizaveta Solovyeva, Moscow Institute of Physics and Technology and INEPCP FRCCP RAS, Moscow, Russia

Marc Vaudel, University of Bergen, Norway

Stefan Schulze, University of Pennsylvania, USA

Veit Schwämmle, University of Southern Denmark, Denmark

Wilhelm, Mathias, Technical University of Munich, Germany

Samaras Patroklos, Technical University of Munich, Germany

Enrique Audaín, University Hospital of Schleswig-Holstein, Kiel, Germany

Juan Antonio Vizcaíno, EMBL-EBI, UK

Melanie Christine Föll, Northeastern University, USA

Pablo Moreno, EMBL-EBI, UK

Johannes Griss, Medical University of Vienna, Austria

Mikhail V. Gorshkov, INEPCP FRCCP RAS, Moscow, Russia

David Bouyssié, IPBS, University of Toulouse, CNRS, UPS, Toulouse, France

Tim Van Den Bossche, Ghent University, Ghent, Belgium

Henry Webel, Novo Nordisk Foundation Center for Protein Research, University of Copenhagen

Julian Uszkoreit, Ruhr University Bochum, Bochum, Germany

Eric W. Deutsch, Institute for Systems Biology

Nuno Bandeira, University of California San Diego

Mingze Bai, Chongqing University of Posts and Telecommunications, Chongqing, China

Lev Levitsky, INEPCP FRCCP RAS, Moscow, Russia

Timo Sachsenberg, Tübingen University, Germany

Yasset Perez-Riverol, EMBL-EBI, UK

Status of this document

This document provides information to the proteomics community about a proposed standard for sample metadata annotations (for instance, to be used in public repositories) called Sample and Data Relationship File (SDRF), as part of the MAGE-TAB Proteomics format. Distribution is unlimited.

Version Draft - this is a draft of version 1.0

### Abstract

The Human Proteome Organisation (HUPO) Proteomics Standards Initiative (PSI) defines community standards for data representation in proteomics to facilitate data comparison, exchange, and verification. This document presents a specification for a sample metadata annotation of proteomics experiments.

Further detailed information, including any updates to this document, implementations, and examples is available at <https://github.com/bigbio/proteomics-metadata-standard>. The official PSI web page for the document is the following [http://psidev.info/](http://psidev.info/sdrf)magetab.

### 1. Introduction

#### **1.1 Description of the need**

Many resources have emerged that provide raw and/or processed proteomics data in the public domain. If these are valuable individually, their integration through re-analysis represents a huge asset for the community [1]. Unfortunately, proteomics experimental design and sample-related information are often missing in public repositories or stored in very diverse ways and formats. For example, the CPTAC consortium (<https://cptac-data-portal.georgetown.edu/>) provides for every dataset a set of excel files with the information on each sample (e.g. <https://cptac-data-portal.georgetown.edu/study-summary/S048>) including tumor size, origin, but also how every sample is related to a specific raw file (e.g. instrument configuration parameters). As a resource routinely re-analyzing public datasets, ProteomicsDB captures for each sample in the database a minimum number of properties to describe the sample and the related experimental protocol such as tissue, digestion method, and instrument (e.g., <https://www.proteomicsdb.org/#projects/4267/6228>). Such heterogeneity often prevents data interpretation, reproducibility, and integration of data from different resources. This is why we propose a data standard for proteomics metadata annotation. For every proteomics dataset we propose to capture at least three levels of metadata: (i) dataset description, (ii) the sample to data files related information; and (iii) the technical/proteomics specific information, available in standard data file formats (e.g., the PSI formats mzIdentML, mzML, or mzTab, among others).

The general dataset description includes minimum information to describe the study overall: title, description, date of publication, type of experiment (e.g., <http://proteomecentral.proteomexchange.org/cgi/GetDataset?ID=PXD016060.0-1&outputMode=XML>). The proteomics standard data files contain mostly the technical metadata associated with the dataset including search engine settings, scores, workflows, configuration files, but do include very limited information about the sample metadata and/or the experimental design*.* Currently, all ProteomeXchange partners mandate both types of information for each dataset. However, the information regarding the samples and its relationship to the data files (Figure 1) is mostly missing [1].

These three levels of metadata are combined in the well-established data formats ISA-TAB [2] (<https://www.isacommons.org/>) or MAGE-TAB [3], which are used in other omics fields such as metabolomics and transcriptomics. In both data formats, a tab-delimited file is used to annotate the sample metadata and link it to the corresponding data file(s). In MAGE-TAB this information is encoded in the sample to data relationship file format - SDRF). Both data formats encode the properties and sample attributes as columns, and each row represents a sample in the study. However, more important than the file format itself, general guidelines about what information needs to be encoded to enable reproducibility of proteomics results is needed. The lack of guidelines to annotate information such as disease stage, cell line code, organism part, or the analytical information about labeling channels (e.g., TMT, SILAC) makes the data representation incomplete. The consequence is that it is not possible to understand the original experiment, and/or perform a re-analysis of the dataset, having all the necessary information for reproducibility purposes. For instance, if the information about the fractions, labeling channels, or enrichment methods is not annotated, the reuse and reproduction of the original results will be challenging, if possible, at all.

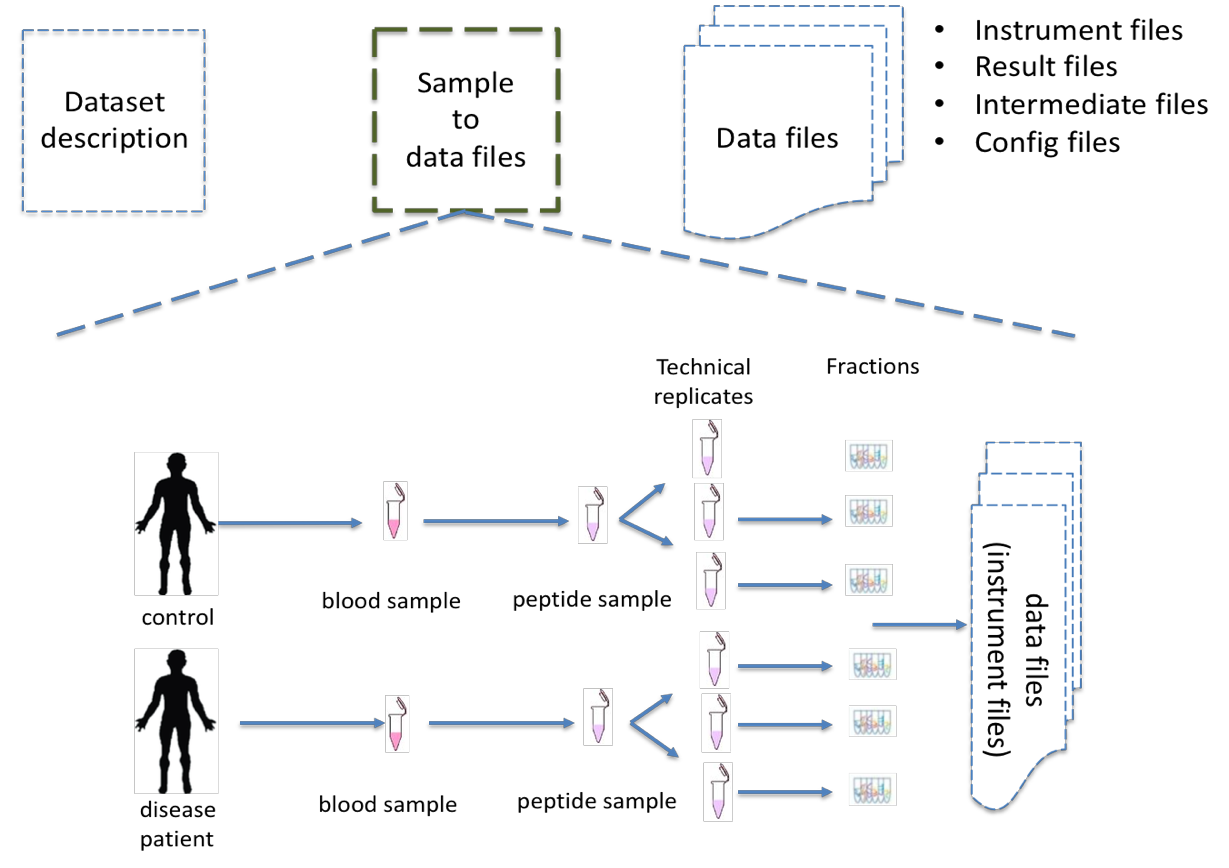

**Figure 1**: The MAGE-TAB-Proteomics file format stores the information of the sample and its relationship to the data files in the dataset. The file format includes not only information about the sample but also about how the data was acquired and processed.

We proposed in this specification to extend the MAGE-TAB file format (<http://fged.org/projects/mage-tab/>) for proteomics data representation. While MAGE-TAB was originally developed for microarray data representation, it has been extended for RNA-Seq and more recently for single-cell transcriptomics experiments [4, 5]. Thousands of transcriptomics datasets have been annotated using MAGE-TAB. By repurposing and extending MAGE-TAB for Proteomics, we aim to provide a format that facilitates the integration of proteomics datasets with other omics data types, i.e., enabling a better representation and linking between multiomics datasets available in different data repositories, including the PX ones.

The MAGE-TAB is divided into two main files, which are linked: IDF (Investigation Description Format) and SDRF (Sample and Data Relationship Format). We will describe how these two files are adapted for Proteomics.

- IDF: The IDF file contains fields describing the study, including e.g., authors/submitters, protocols, publications. ProteomerXchange resources have developed a file format called submission.px which captures equivalent information to MAGE-TAB IDF. We have developed a set of tools to automatically convert from submission.px to IDF (<https://github.com/bigbio/proteomics-metadata-standard/blob/master/generate_idf.py>).
- SDRF: The Sample to Data files relationship information is missing (or not standardized) for all PX datasets.

The original IDF file format developed in MAGE-TAB does not need to change to represent proteomics data. The standardization of the SDRF for proteomics is the main objective of this specification document.

#### **1.2 Requirements of SDRF-Proteomics**

The MAGE-TAB-Proteomics is based on the MAGE-TAB file format describes the sample characteristics and the relationships between samples and data files included in a dataset. The information in SDRF files is organized so that it follows the natural flow of a proteomics experiment. The main requirements to be fulfilled for the SDRF-Proteomics format are:

- The SDRF file is a tab-delimited format where each row corresponds to a relationship between a Sample and a Data file.
- Each column MUST correspond to an attribute/property of the Sample or the Data file.
- Each value in each cell MUST be the property for a given Sample or Data file.
- The file MUST begin with columns describing the samples and continue the data files generated from the analyses of the experimental results.
- Support for handling unknown values/characteristics.
- The SDRF-Proteomics does not aim to capture the downstream analysis part of the experimental design including details of which samples were compared to which other samples, or how samples are combined into study variables or parameters for the downstream analysis such as FDR or p-values thresholds. The HUPO-PSI community will work in the future to include this information in other file formats such as mzTab or a new type of file format.

#### **1.3 Issues to be addressed**

The main issues to be addressed by the SDRF are:

- It MUST be able to represent the sample metadata and the metadata of the data files generated by the instruments or the analyses.
- It MUST be able to represent the experimental design including the way samples and data have been collected.

### 2. Notational Conventions

The keywords “MUST”, “MUST NOT”, “REQUIRED”, “SHALL”, “SHALL NOT”, “SHOULD”, “SHOULD NOT”, “RECOMMEND/RECOMMENDED”, “MAY”, “COULD BE”, and “OPTIONAL” are to be interpreted as described in RFC 2119 (2).

### 3. Documentation

The official website for the SDRF-Proteomics project is <https://github.com/bigbio/proteomics-metadata-standard>. New use cases, changes to the specification, and examples can be added by using Pull requests or issues in GitHub (see the introduction to GitHub - <https://lab.github.com/githubtraining/introduction-to-github>).

A set of examples and annotated projects from ProteomeXchange can be found here: <https://github.com/bigbio/proteomics-metadata-standard/tree/master/annotated-projects>

Multiple tools have been implemented to validate SDRF-Proteomics files:

- sdrf-pipelines (Python - <https://github.com/bigbio/sdrf-pipelines>): This tool allows to validation of an SDRF-Proteomics file. In addition, allows converting SDRF to other popular pipelines and software configuration files such as MaxQuant or OpenMS.

- jsdrf (Java - <https://github.com/bigbio/jsdrf>): This Java library and tool allows to validate SDRF-Proteomics files. It also includes a generic data model that can be used by Java applications.

### 4. Relationship to other specifications

As mentioned above, SDRF-Proteomics is fully compatible with the SDRF file format part of MAGE-TAB (<https://www.ebi.ac.uk/arrayexpress/help/magetab_spec.html>). The MAGE-TAB was originally developed to store the metadata and sample information on microarray experiments and has been extended to support RNA-Seq and single-cell RNA-Seq experiments. If a ProteomeXchange px project file is converted to an IDF file (project description in MAGE-TAB) and is combined with the SDRF-Proteomics, a valid MAGE-TAB Proteomics is obtained.

SDRF-Proteomics sample information can also be embedded into mzTab files ([https://www.psidev.info/mztab).](https://www.psidev.info/mztab) The sample metadata in mzTab contains properties such as the column names in the SDRF-Proteomics and values as sample cell values.

SDRF-Proteomics aims to capture the sample metadata and its relationship with the data files (e.g. raw files from mass spectrometers).

### 5. Ontologies

The list of ontologies/controlled vocabularies (CV) supported in SDRF-Proteomics are:

- PSI Mass Spectrometry CV ([PSI-MS](https://www.ebi.ac.uk/ols/ontologies/ms))
- Experimental Factor Ontology ([EFO](https://www.ebi.ac.uk/ols/ontologies/efo)).
- [Unimod](https://www.ebi.ac.uk/ols/ontologies/unimod) protein modification database for mass spectrometry
- PSI-MOD CV ([PSI-MOD](https://www.ebi.ac.uk/ols/ontologies/mod))
- [Cell line ontology](https://www.ebi.ac.uk/ols/ontologies/clo)
- [Drosophila anatomy ontology](https://www.ebi.ac.uk/ols/ontologies/fbbt)
- [Cell ontology](https://www.ebi.ac.uk/ols/ontologies/cl)
- [Plant ontology](https://www.ebi.ac.uk/ols/ontologies/po)
- [Uber-anatomy ontology](https://www.ebi.ac.uk/ols/ontologies/uberon)
- [Zebrafish anatomy and development ontology](https://www.ebi.ac.uk/ols/ontologies/zfa)
- [Zebrafish developmental stages ontology](https://www.ebi.ac.uk/ols/ontologies/zfs)
- [FlyBase Developmental Ontology](https://www.ebi.ac.uk/ols/ontologies/fbdv)
- [Rat Strain Ontology](https://www.ebi.ac.uk/ols/ontologies/rs)
- [Chemical Entities of Biological Interest Ontology](https://www.ebi.ac.uk/ols/ontologies/chebi)
- [NCBI organismal classification](https://www.ebi.ac.uk/ols/ontologies/ncbitaxon)
- [PATO - the Phenotype and Trait Ontology](https://www.ebi.ac.uk/ols/ontologies/pato)
- PRIDE Controlled Vocabulary ([PRIDE](https://www.ebi.ac.uk/ols/ontologies/pride))

If an additional ontology/CV not included in the previous list is needed for encoding a specific characteristic or comment, it is possible to do it. However, the ontology/CV SHOULD be accessible through the OLS service (<https://www.ebi.ac.uk/ols/index>).

### 6. SDRF-Proteomics file format

The SDRF-Proteomics file format describes the sample characteristics and the relationships between samples and data files. The file format is a **tab-delimited** one where each row encodes a relationship between a Sample and a Data file, and each column corresponds to an attribute/property of the Sample or of the Data file, and the value in each cell is the specific value of the property for a given Sample (**Figure 2**).

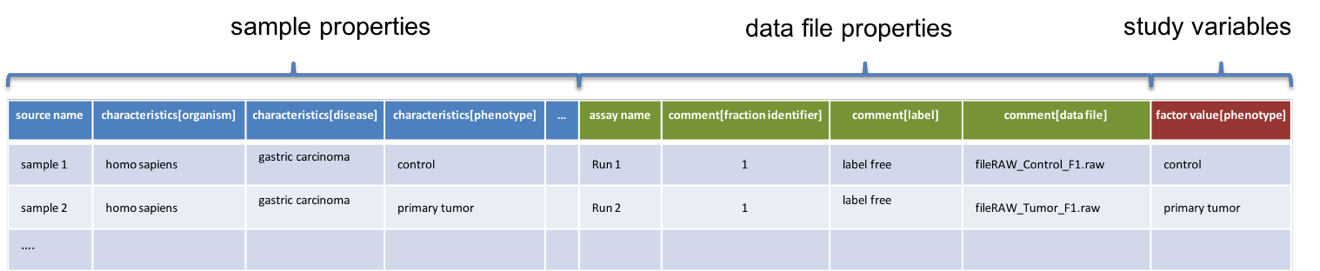

**Figure 2**: SDRF-Proteomics in a nutshell. The file format is a tab-delimited one where **columns** encode properties of the samples, the data files, or the variables under study. The **rows** are Sample/Data file relationships and the **cells** are the given values for one property in a specific sample.

#### **6.1 Format rules**

There are general scenarios/use cases that are addressed by the following rules:

- **Unknown values**: In some cases, the column is mandatory in the format but for some samples the corresponding value is unknown. In those cases, users SHOULD use ‘not available’ (if the value is not known).
- **Not Applicable values**: In some cases, the column is mandatory but for some samples, the corresponding value is not applicable. In those cases, users SHOULD use ‘not applicable’.
- **Case sensitivity**: By specification, the SDRF is case insensitive, but we RECOMMEND using lowercase characters throughout all the text (Column names and values).
- **Spaces**: By specification, SDRF is space sensitive (sourcename != source name).
- **Column order**: SDRF MUST start with the **source name** column (accession/name of the sample), then all the sample characteristics, followed by the **assay name**. Finally, after the assay name, all the comments (properties of the data file generated) MUST be included.
- **Extension**: The extension of the SDRF format should be .tsv or .txt.

#### **6.2 SDRF values**

The value for each property (e.g., *characteristics*, *comment*) corresponding to each sample can be represented in multiple ways.

A) Free Text (Human readable): In the free text representation, the value is provided as text without ontology/CV support (e.g., colon or providing accession numbers). This is only RECOMMENDED when the **text** inserted in the table is the exact *name* of an ontology/CV term in EFO. If the term is not included in EFO, other ontologies can be used.

| **source name** | **characteristics[organism]** |
| --- | --- |
| sample 1 | homo sapiens |
| sample 2 | homo sapiens |

B) Ontology URL (Computer-readable): Users can provide the corresponding ontology/CV accession number for one value. This is recommended for enriched files where the user does not want to use intermediate tools to map from free text to ontology/CV terms.

| **source name** | **characteristics[organism]** |
| --- | --- |
| Sample 1 | [NCBITaxon:9606](http://purl.obolibrary.org/obo/NCBITaxon_9606) |
| Sample 2 | [NCBITaxon:9606](http://purl.obolibrary.org/obo/NCBITaxon_9606) |

C) Key=value representation (Human and Computer-readable): The current representation aims to provide a mechanism to represent the complete information of the ontology/CV term including *Accession*, *Name*, and other additional properties.

In the key=value pair representation, the value of the property is represented as an object with multiple properties, where the key is one of the properties of the object and the value is the corresponding value for the particular key. For example (for protein modifications – **section 6.5.1**):

NT=Glu->pyro-Glu; MT=fixed; PP=Anywhere; AC=Unimod:27; TA=E

### 7. Minimum information about Samples

The Sample metadata has different **Categories/Headings** to organize all the attributes/column headers available of a given sample. Each Sample in the dataset MUST contain a *source name* and a collection of *characteristics*:

| Property | Mandatory (1)  Optional (0) | Cardinality | Description | Example |
| --- | --- | --- | --- | --- |
| source  name | 1 | 1 | Unique sample name (it can be present multiple times if the same sample is used several times in the same dataset) | Sample 1 |
| characteristics | 1 | 1.. * | “characteristics” column headings SHOULD contain an ontology property term in square brackets. Multiple *Characteristic* columns of the same category (e.g., “characteristics [organism part]”) are allowed. Typically, the usage implies whole to part from left to right. | characteristics [Organism Part] |

Example:

| source name | characteristics[organism] | characteristics[phenotype] | characteristics[compound] |
| --- | --- | --- | --- |
| Sample1 | homo sapiens | necrotic tissue | drug A |
| Sample2 | homo sapiens | normal | none |

Some important notes:

Each *characteristics* name in the column header SHOULD be a CV term from the [EFO ontology](https://www.ebi.ac.uk/ols/ontologies/efo). For example, the Header *characteristics[organism]* correspond to the ontology term [Organism](http://www.ebi.ac.uk/efo/EFO_0000634).

Multiple values (columns) for the same *characteristics* term are allowed in SDRF-Proteomics. However, it is RECOMMENDED not to use the same column in the same SDRF. If you have multiple phenotypes, you can specify what it refers to or use another more specific term, e.g. "immunophenotype".

### 8. From Samples to Data files

The connection between the *Samples* to the Data files is done by using a series of properties and attributes. All the properties referring to the MS run (file) itself are annotated with the category ***comment****.* The use of *comment* is mainly aimed at differentiating sample properties from the data file properties. It matches a given *sample* to the corresponding *file(s).* The word ***comment*** is used for backward-compatibility with gene expression experiments (RNA-Seq and Microarrays experiments) since this is the mechanism used in the original SDRF for transcriptomics.

The order of the columns in the file format is important: ***assay name*** MUST always be located before the comments. It is RECOMMENDED to put the last column as a *comment[data file]* before the next category called **factor values**.

The following properties SHOULD be provided for each data file (MSRun) file:

- *assay name*: For SDRF backward compatibility MSRun cannot be used. Instead, the assay *name* is used. Examples of assay names are: “run 1”, “run_fraction_1_2”. The name of the file can be used as an assay name.
- *comment[fraction identifier]*: The *fraction identifier* allows recording the number of a given fraction. The fraction identifier corresponds to this [ontology term](https://www.ebi.ac.uk/ols/ontologies/ms/terms?iri=http%3A%2F%2Fpurl.obolibrary.org%2Fobo%2FMS_1000858). It MUST start from **1** and if the experiment does not contain fractions, “1” MUST be used for each MSRun (*assay*).
- *comment[label]*: it describes the label applied to each Sample (if any). In case of multiplex experiments such as TMT, SILAC, and/or ITRAQ the corresponding *label* SHOULD be added. For label-free experiments, the *label-free sample* term MUST be used. The terms that SHOULD be used in this column are in the PRIDE Ontology under the [Label category](https://www.ebi.ac.uk/ols/ontologies/pride/terms?iri=http%3A%2F%2Fpurl.obolibrary.org%2Fobo%2FPRIDE_0000514&viewMode=All&siblings=false).
- *comment[data file]*: The *data file* provides the name of the raw file generated by the instrument. The data files can be instrument raw files but also converted peak lists such as mzML, MGF.

|  | assay name | comment[label] | comment[fraction identifier] | comment[data file] |
| --- | --- | --- | --- | --- |
| sample 1 | run 1 | label free sample | 1 | 000261_C05_P0001563_A00_B00K_R1.RAW |
| sample 1 | run 2 | label free sample | 2 | 000261_C05_P0001563_A00_B00K_R2.RAW |

All the possible *label* values can be seen in the PRIDE CV under the [Label](https://www.ebi.ac.uk/ols/ontologies/pride/terms?iri=http%3A%2F%2Fpurl.obolibrary.org%2Fobo%2FPRIDE_0000514&viewMode=All&siblings=false) node.

#### **8.1 Encoding sample technical and biological replicates**

Different measurements of the same biological sample are often categorized as (i) *Technical* or (ii) *Biological* replicates, based on whether they are (i) matched on all variables, e.g., same sample and same protocol; or (ii) different samples matched on study variable(s), e.g., different patients receiving a placebo, or in placebo vs. drug trial. Technical and biological replicates have different levels of independence, which must be taken into account during data interpretation.

For a given experiment, there are different levels to which samples can be matched - e.g., same sample, sample protocol, covariates - the definition of technical replicate can therefore vary based on the number of variables included. The technical and biological replicates in SDRF depend on the factor variable under study. For example, if the aim is to quantify protein expression among different tissues probably the biological replicates are different than if the study would aim to compare expression considering tumor size in different tissues.

**Technical replicate**: It is defined as repeated measurements of the same sample, that represent independent measures of both signal and noise associated with protocols or equipment [6]. In the SDRF-Proteomics, technical replicate information MUST be provided *comment[technical replicate]*.

In MS-based proteomics a technical replicate can be, for example, doing the full sample preparation from extraction to MS multiple times to control variability in the instrument and sample preparation. Another valid example would be to replicate only one part of the analytical method, for example, run the sample twice on the LC-MS/MS. Technical replicates indicate if measurements are scientifically robust or noisy, and how large the measured effect must be to stand out above that noise.

In the following example, the technical replicate column enables to distinguish quantitative values of the same fraction but obtained from different technical replicates.

| source name | assay name | comment[label] | comment[fraction identifier] | comment[technical replicate] | comment[data file] |
| --- | --- | --- | --- | --- | --- |
| Sample 1 | run 1 | label-free sample | 1 | 1 | 000261_C05_P0001563_A00_B00K_F1_TR1.RAW |
| Sample 1 | run 2 | label free sample | 2 | 1 | 000261_C05_P0001563_A00_B00K_F2_TR1.RAW |
| Sample 1 | run 3 | label free sample | 1 | 2 | 000261_C05_P0001563_A00_B00K_F1_TR2.RAW |
| Sample 1 | run 4 | label free sample | 2 | 2 | 000261_C05_P0001563_A00_B00K_F2_TR2.RAW |

In cases where a single sample is analyzed in multiple different ionization modes (e.g. positive and negative), these are considered neither technical replicates nor fractions, but rather different experimental conditions.

**Biological replicate**: It is defined as independent measurements of biologically distinct samples that capture biological variation, which may itself be a subject of study or a source of variation unrelated to the purposes of the study. In SDRF-Proteomics, samples with the same characteristics used for the same factor values are considered biological replicates. Biological replicates MUST be annotated using *characteristics[biological replicate]*.

One example containing explicit annotation of the biological replicates can be found here:

<https://github.com/bigbio/proteomics-metadata-standard/blob/c3a56b076ef381280dfcb0140d2520126ace53ff/annotated-projects/PXD006401/sdrf.tsv>

### 9. Data properties

#### **9.1 Type and Model of Mass Spectrometer**

The mass spectrometer model SHOULD be specified as a *comment[instrument]*. Possible values are listed under the [instrument model](https://www.ebi.ac.uk/ols/ontologies/ms/terms?iri=http%3A%2F%2Fpurl.obolibrary.org%2Fobo%2FMS_1000031&viewMode=All&siblings=false) term. Additionally, it is strongly RECOMMENDED to include *comment[MS2 analyzer type]*. This is important e.g., for Orbitrap models where MS2 scans can be acquired either in the Orbitrap or in the ion trap. Setting this value allows differentiating high-resolution MS/MS data. Possible values of *comment[MS2 analyzer type]* are the following [mass analyzer types](https://www.ebi.ac.uk/ols/ontologies/ms/terms?iri=http%3A%2F%2Fpurl.obolibrary.org%2Fobo%2FMS_1000443&viewMode=All&siblings=false).

#### **9.2 Label annotations**

In order to annotate quantitative datasets, SDRF makes use of tags for each channel associated with the sample in a column named *comment[label]*. The label values are organized under the following ontology term [Label](https://www.ebi.ac.uk/ols/ontologies/pride/terms?iri=http%3A%2F%2Fpurl.obolibrary.org%2Fobo%2FPRIDE_0000514&viewMode=All&siblings=false). Some of the most popular labels are:

- For label-free experiments, the value SHOULD be “label free sample”.
- For TMT experiments the SDRF can use the PRIDE ontology terms under the sample label. Here some examples of TMT channels:
  - TMT126, TMT127, TMT127C, TMT127N, TMT128, TMT128C, TMT128N, TMT129, TMT129C, TMT129N, TMT130, TMT130C, TMT130N, TMT131

In order to achieve a clear relationship between the label and the sample characteristics, each channel of each sample (in multiplex experiments) MUST be defined in a separate row: **one row per channel used (annotated with the corresponding *comment[label]* per file**.

#### **9.3 Additional RAW file properties**

It is RECOMMENDED to include the public URI of the files, if available. For example, for ProteomeXchange datasets, the URI from the FTP location COULD be provided:

| source name | **…** | **comment[file uri]** |
| --- | --- | --- |
| sample 1 | … | ftp://ftp.pride.ebi.ac.uk/pride/data/archive/2017/09/PXD005946/000261_C05_P0001563_A00_B00K_R1.RAW |

#### **9.4 Data file additional properties**

This section includes some additional properties that have been agreed on so that they RECOMMEND be captured for each proteomics experiment.

- *comment[fractionation method]*: The fraction method used to separate the sample. The values of this term can be read under the PRIDE ontology term [Fractionation method](https://www.ebi.ac.uk/ols/ontologies/pride/terms?iri=http%3A%2F%2Fpurl.obolibrary.org%2Fobo%2FPRIDE_0000550). Example: “Off-gel electrophoresis”.
- *comment[depletion]*: The removal of specific components of a complex mixture of proteins or peptides based on some specific property of those components. The values of the column will be “no depletion” or “depletion”.
- *comment[collision energy]*: Collision energy COULD added as non-normalized (10000 eV) or normalized (1000 NCE) value.
- *comment[dissociation method]*: This property COULD include information about the fragmentation method, like e.g. HCD, CID. The values of the column are under the term [dissociation method](https://www.ebi.ac.uk/ols/ontologies/ms/terms?iri=http%3A%2F%2Fpurl.obolibrary.org%2Fobo%2FMS_1000044&viewMode=All&siblings=false).

#### **9.5 Data technical details properties**

It is RECOMMENDED to encode some of the technical parameters of the MS experiment as comments including the following parameters:

- Protein Modifications
- Precursor and fragment ion mass tolerances
- Digestion Enzymes

##### **9.5.1 Protein Modifications**

Sample modifications (including both chemical modifications and post-translational modifications, PTMs) are originated from multiple sources: **artifactual modifications**, **isotope labeling**, adducts that are encoded as PTMs (e.g., sodium) or the most **biologically relevant** PTMs.

It is RECOMMENDED to provide the modifications expected (they are encoded as modification parameters) in the sample including the amino acids affected, whether they are Variable or Fixed (also Custom and Annotated modifications are supported).

The RECOMMENDED name of the column for sample modification parameters is a *comment[modification parameters]*. Each modification parameter represents a unique value of a PTM in the SDRF-Proteomics. If multiple modifications are used in the experiment, multiple columns (*comment[modification parameters]*) MUST be used.

The modification parameters correspond to the name of the ontology term [MS:1001055](https://www.ebi.ac.uk/ols/ontologies/ms/terms?iri=http%3A%2F%2Fpurl.obolibrary.org%2Fobo%2FMS_1001055).

For each modification, different properties are captured using a key=value pair structure, including name, position, etc. All the possible features available for modification parameters are:

| **Property** | **Key** | **Example** | **Mandatory(1)**  **Optional(0️)** | **comment** |
| --- | --- | --- | --- | --- |
| Name of the Modification | NT | NT=Acetylation | 1 | Name of the Term in this particular case. In the case of custom modifications, it can be a name defined by the user. |
| Modification Accession | AC | AC=UNIMOD:1 | 0️ | Accession number in an external database. UNIMOD or PSI-MOD are currently supported. |
| Chemical Formula | CF | CF=H(2)C(2)O | 0️ | This is the chemical formula of the added or removed atoms. For the formula composition please follow the guidelines from [UNIMOD](http://www.unimod.org/names.html). |
| Modification Type | MT | MT=Fixed | 0️ | This specifies which modification group the modification belongs to. The choices are: [Fixed, Variable, Annotated]. The term *Annotated* is used to enable the search for all the occurrences of the modification into an annotated protein database file such as UNIPROT XML or PEFF. |
| Position of the modification in the Polypeptide | PP | PP=Any N-term | 0️ | Choose from the following options: [Anywhere, Protein N-term, Protein C-term, Any N-term, Any C-term]. The default is **Anywhere**. |
| Target Amino acid | TA | TA=S,T,Y | 1 | The target amino acid residue. If the modification targets multiple sites, it can be separated by the TA (see below). |
| Monoisotopic Mass | MM | MM=42.010565 | 0️ | The exact monoisotopic atomic mass shift produced by the modification. Please use at least 5 decimal places of accuracy. This SHOULD only be used if the chemical formula of the modification is not known. If the chemical formula is specified, the monoisotopic mass will be overwritten by the calculated monoisotopic mass. |
| Target Site | TS | TS=N[^P][ST] | 0️ | For some software, it is important to capture complex rules for modification sites as regular expressions. These use cases SHOULD be specified as regular expressions. |

We RECOMMEND using the Unimod interim name or the PSI-MOD name for indicating the modification name. For custom modifications, we RECOMMEND using an intuitive name. If the PTM is unknown (custom), the *Chemical Formula* or *Monoisotopic Mass* MUST be annotated.

An example of an SDRF-Proteomics file with sample modifications annotated, where each modification needs an extra column in the file format:

| source name | comment[modification parameters] | comment[modification parameters] |
| --- | --- | --- |
| sample 1 | NT=Glu-pyro-Glu; MT=fixed; PP=Anywhere; AC=Unimod:27; TA=E | NT=Oxidation; MT=Variable; TA=M |

##### **9.5.2 Cleavage agents**

The REQUIRED *comment [cleavage agent details]* property is used to capture the enzyme information. The list of Enzyme names can be found in the PSI-MS CV (<https://www.ebi.ac.uk/ols/ontologies/ms/terms?iri=http%3A%2F%2Fpurl.obolibrary.org%2Fobo%2FMS_1001045&viewMode=All&siblings=false>). Similarly to protein modifications, a key=value pair representation is used to encode the following properties for each enzyme:

| **Property** | **Key** | **Example** | **Mandatory(1)**  **Optional(0️)** | **comment** |
| --- | --- | --- | --- | --- |
| Name of the Enzyme | NT | NT=Trypsin | 1 | Name of the Enzyme. |
| Enzyme Accession | AC | AC=MS:1001251 | 0️ | Accession in the external PSI-MS CV definition under the following category [Cleavage agent name](https://www.ebi.ac.uk/ols/ontologies/ms/terms?iri=http%3A%2F%2Fpurl.obolibrary.org%2Fobo%2FMS_1001045). |
| Cleavage site regular expression | CS | CS=(?⇐[KR])(?!P) | 0️ | The cleavage site is defined as a regular expression. |

An example of an SDRF-Proteomics section with the sample enzyme annotated:

| source name | comment[cleavage agent details] |
| --- | --- |
| sample 1 | NT=Trypsin; AC=MS:1001251; CS=(?⇐[KR])(?!P) |

If no enzyme is used, for example in the case of top-down experiments or (some) peptidomics experiments, the value SHOULD be ‘not applicable’.

##### **9.5.3 Precursor and Fragment mass tolerances**

For proteomics experiments, it is important to encode different mass tolerances (for precursor and fragment ions).

| source name | comment[fragment mass tolerance] | comment[precursor mass tolerance] |
| --- | --- | --- |
| sample 1 | 0.6 Da | 20 ppm |

Units for the mass tolerances (either Da or ppm) MUST be provided. A unique value should be provided by each RAW file. For example, in multiplex experiments, multiple samples would be in multiple rows associated with the same (assay) RAW file.

### 10. SDRF-Proteomics study variables

The variable/property under study MUST be highlighted using the **factor value** category. For example, the *factor value[disease]* is used when the main purpose of a given experiment is to compare protein expression across different diseases or different states of a given disease. Multiple variables under study can be included by adding multiple factor values columns.

| factor value | 1..* | “factor value” columns SHOULD indicate which experimental factor/variable is used as the hypothesis to perform the data analysis. The “factor value” columns SHOULD occur after all characteristics and attributes of the samples. | Factor Value [phenotype] |
| --- | --- | --- | --- |

### 11. Specific use cases and conventions

Conventions define how to encode some particular information in the file format by supporting specific use cases. Conventions define a set of new columns that are needed to represent a particular use case or experiment type (e.g. phosphorylation-enriched dataset). In addition, conventions define how some specific free-text columns (values that are not defined as ontology terms) should be written.

Conventions are documented and compiled from at <https://github.com/bigbio/proteomics-metadata-standard/issues> or by performing a pull-request. New conventions will be added to updated versions of this specification document in the future. It is planned that, unlike in other PSI formats, more regular updates will need to be done to be able to explain how new use cases for the format can be accommodated.

#### **11.1 How to encode age and other elapsed times**

One of the characteristics of a sample can be the age of an individual. It is RECOMMENDED to provide the age in the following format: {X}Y{X}M{X}D. Some valid examples are:

- 40Y (forty years)
- 40Y5M (forty years and 5 months)
- 40Y5M2D (forty years, 5 months, and 2 days)

When needed, weeks can also be used: 8W (eight weeks)

Age interval:

Sometimes the sample does not have an exact age but contains a range of ages. To annotate an age range the following convention is RECOMMENDED:

40Y-85Y

This means that the subject (sample) is between 40 and 85 years old.

Other temporal information can be encoded similarly.

#### **11.2 Phosphoproteomics and other post-translational modifications enriched studies**

In PTM-enriched experiments, the *characteristics[enrichment process]* SHOULD be provided. The different values already included in EFO are:

- enrichment of phosphorylated proteins
- enrichment of glycosylated proteins

This characteristic can be used as a *factor value[enrichment process]* to differentiate the expression between proteins in the phospho-enriched sample when compared with the control.

#### **11.3 Pooled samples**

When multiple samples are pooled into one, the general approach is to annotate them separately, abiding by the general rule: one row stands for one sample-to-one-file relationship. In this case, multiple rows are created for the same corresponding data file.

One exception is made for the case when one channel e.g., in a TMT/iTRAQ multiplexed experiment, is used for a sample pooled from other channels, typically for normalization purposes. In this case, it is not necessary to repeat all sample annotations. Instead, a special characteristic value can be used:

| **source name** | **characteristics[pooled sample]** | **assay name** | **comment[label]** | **comment[data file]** |
| --- | --- | --- | --- | --- |
| sample 1 | not pooled | run 1 | TMT131 | file01.raw |
| sample 2 | not pooled | run 1 | TMT131C | file01.raw |
| sample 10 | SN=sample 1,sample 2, … sample 9 | run 1 | TMT128 | file01.raw |

SN stands for source names and can be used to list source name fields of samples that are annotated in the same file and that are **used in the same experiment and the same MS run**.

Another possible value for *characteristics[pooled sample]* is the string “pooled” for cases when it is known that a sample is pooled but the individual samples included in that pool cannot be annotated.

#### **11.4 Synthetic peptide libraries**

It is common to use synthetic peptide libraries for multiple use cases including:

- Benchmark of analytical and bioinformatics methods and algorithms.
- Improvement of peptide identification/quantification using spectral libraries.

When describing synthetic peptide libraries most of the sample metadata can be declared as “not applicable”. However, some authors can also annotate the organism, for example, because they know that the library has been designed from specific peptide species, see example the following experiment containing synthetic peptides (https://github.com/bigbio/proteomics-metadata-standard/blob/master/annotated-projects/PXD000759/PXD000759.sdrf.tsv).

In these cases, it is important to annotate that the sample is composed of a synthetic peptide library. This can be done by adding the *characteristics[synthetic peptide]*. The possible values are “synthetic”,  “not synthetic” or “mixed”.

#### **11.5 Normal and healthy samples**

Samples from healthy patients or individuals normally appear in manuscripts and are often annotated as healthy or normal. We RECOMMEND using the word “normal” mapped to the CV term PATO_0000461, which is also included in EFO: [normal PATO term](https://www.ebi.ac.uk/ols/ontologies/efo/terms?iri=http%3A%2F%2Fpurl.obolibrary.org%2Fobo%2FPATO_0000461). Example:

| **source name** | **characteristics[organism]** | **characteristics [organism part]** | **characteristics[phenotype]** | **characteristics[compound]** | **factor value[phenotype]** |
| --- | --- | --- | --- | --- | --- |
| sample_treat | homo sapiens | liver | necrotic tissue | drug A | necrotic tissue |
| sample_control | homo sapiens | liver | normal | none | normal |

#### **11.6 Multiple projects included into one SDRF file**

It may be needed to annotate multiple ProteomeXchange datasets into one large SDRF-Proteomics file e.g., reanalysis purposes. If that is the case, it is RECOMMENDED to use the column name *comment[proteomexchange accession number]* to differentiate between different datasets.

### 12. SDRF-Proteomics templates

The **sample metadata templates** contain the set of guidelines needed to annotate different types of proteomics experiments to ensure that minimum metadata and characteristics are provided to be able to understand the experimental design of a dataset. These templates correspond to the distribution and frequency of experiment types in public databases like [PRIDE](http://www.ebi.ac.uk/pride/archive) and others included in [ProteomeXchange](http://www.proteomexchange.org/):

- Default: Minimum information for any proteomics experiment. [Template](https://github.com/bigbio/proteomics-metadata-standard/blob/master/templates/sdrf-default.tsv)
- Human: All tissue-based experiments that use Human samples. [Template](https://github.com/bigbio/proteomics-metadata-standard/blob/master/templates/sdrf-human.tsv)
- Vertebrates: Vertebrate experiment. [Template](https://github.com/bigbio/proteomics-metadata-standard/blob/master/templates/sdrf-vertebrates.tsv)
- Non-vertebrates: Non-vertebrate experiment. [Template](https://github.com/bigbio/proteomics-metadata-standard/blob/master/templates/sdrf-nonvertebrates.tsv)
- Plants: Plant experiment. [Template](https://github.com/bigbio/proteomics-metadata-standard/blob/master/templates/sdrf-plants.tsv)
- Cell lines: Experiments using cell-lines. [Template](https://github.com/bigbio/proteomics-metadata-standard/blob/master/templates/sdrf-cell-line.tsv)

**Sample attributes**: Minimum sample attributes for primary cells coming from different species and cell lines

|  | Default | Human | Vertebrates | Non-vertebrates | Plants | Cell lines | Ontology/CV URL |
| --- | --- | --- | --- | --- | --- | --- | --- |
| Source Name | 1 | 1 | 1 | 1 | 1 | 1 |  |
| characteristics[organism] | 1 | 1 | 1 | 1 | 1 | 1 | <https://www.ebi.ac.uk/ols/ontologies/ncbitaxon> |
| characteristics[strain/breed] | 0 | 0 | 0 | 0️ | 0 | 0️ | <https://www.ebi.ac.uk/ols/ontologies/efo> |
| characteristics[ecotype/cultivar] | 0 | 0 | 0 | 0 | 0️ | 0 | <https://www.ebi.ac.uk/ols/ontologies/efo> |
| characteristics[ancestry category] | 0 | 1 | 0 | 0 | 0 | 0 | <https://www.ebi.ac.uk/ols/ontologies/efo> |
| characteristics[age] | 0 | 1 | 0️ | 0 | 0️ | 0 |  |
| characteristics[developmental stage] | 0 | 0️ | 0️ | 0 | 0️ | 0 | <https://www.ebi.ac.uk/ols/ontologies/efo> |
| characteristics[sex] | 0 | 1 | 0️ | 0 | 0 | 0 | <https://www.ebi.ac.uk/ols/ontologies/efo> |
| characteristics[disease] | 1 | 1 | 1 | 1 | 0 | 1 | <https://www.ebi.ac.uk/ols/ontologies/efo> |
| characteristics[organism part] | 1 | 1 | 1 | 1 | 1 | 1 | <https://www.ebi.ac.uk/ols/ontologies/efo> |
| characteristics[cell type] | 1 | 1 | 1 | 1 | 1 | 1 | <https://www.ebi.ac.uk/ols/ontologies/efo> |
| characteristics[individual] | 0 | 1 | 0️ | 0️ | 0️ | 0️ |  |
| characteristics[cultured cell] | 0 | 0 | 0 | 0 | 0 | 1 | <https://www.ebi.ac.uk/ols/ontologies/efo> |
| characteristics[biological replicate] | 1 | 1 | 1 | 1 | 1 | 1 |  |
| comment[data file] | 1 | 1 | 1 | 1 | 1 | 1 |  |
| comment[fraction identifier] | 1 | 1 | 1 | 1 | 1 | 1 |  |
| comment[label] | 1 | 1 | 1 | 1 | 1 | 1 | <https://www.ebi.ac.uk/ols/ontologies/ms>  <https://www.ebi.ac.uk/ols/ontologies/pride> |
| comment[cleavage agent details] | 1 | 1 | 1 | 1 | 1 | 1 | <https://www.ebi.ac.uk/ols/ontologies/ms> |
| comment[instrument] | 1 | 1 | 1 | 1 | 1 | 1 | <https://www.ebi.ac.uk/ols/ontologies/ms> |
| comment[technical replicate] | 1 | 1 | 1 | 1 | 1 | 1 |  |

1: Required Attributes for each sample type (e.g. human, vertebrates).

0: Optional Attributes

### 13. Examples of annotated datasets

| Dataset Type | ProteomeXchange / PubMed Accession | MAGE-TAB | Experiment Comments |
| --- | --- | --- | --- |
| Label-free | PXD008934 | <https://github.com/bigbio/proteomics-metadata-standard/tree/master/annotated-projects/PXD008934> |  |
| TMT | CPTAC PMID27251275 | <https://raw.githubusercontent.com/bigbio/proteomics-metadata-standard/master/annotated-projects/PMID27251275/sdrf.tsv> | CPTAC dataset not in ProteomeXchange |
| SILAC | PXD006877 | <https://raw.githubusercontent.com/bigbio/proteomics-metadata-standard/7de8ea98118fb9f7d2958656ada07bf767a52106/annotated-projects/PXD006877/sdrf.tsv> |  |
| Phospho-proteomics | PXD006482 | <https://raw.githubusercontent.com/bigbio/proteomics-metadata-standard/fd0531411a3287abf6a52fc659f6d32f47c20292/annotated-projects/PXD006482/sdrf.tsv> |  |
| Label-free | PXD010154 | <https://github.com/bigbio/proteomics-metadata-standard/tree/master/annotated-projects/PXD010154> | Profile, 29 tissues dataset, include tonsil acquired with various enzymes and fragmentation modes. |
| AP-MS interactomics | PXD018117 | <https://github.com/bigbio/proteomics-metadata-standard/tree/master/annotated-projects/PXD018117> | SARS-CoV-2, AP-MS, label-free |
|  | PXD005336 |  | Drug concentrations, Clinical Kinase inhibitors |
| TMT | PXD017710 | <https://github.com/bigbio/proteomics-metadata-standard/tree/master/annotated-projects/PXD017710> | time points, SARS-CoV-2 infection time points, TMT |
| Label-free | PXD004242 | <https://github.com/bigbio/proteomics-metadata-standard/tree/master/annotated-projects/PXD004242> | paired samples, weight loss, multiple time points for the same patients |

### 14. Authors Information

Chengxin Dai

Chongqing University of Posts and Telecommunications

Chongqing, China

Juan Antonio Vizcaíno

European Bioinformatics Institute (EMBL-EBI),

Cambridge, United Kingdom

Anja Fullgrabe

European Bioinformatics Institute (EMBL-EBI),

Cambridge, United Kingdom

Mingze Bai

Chongqing University of Posts and Telecommunications

Chongqing, China

Marie Locard-Paulet

Novo Nordisk Foundation Center for Protein Research, University of Copenhagen, Copenhagen, Denmark

Veit Schwämmle

Department of Biochemistry and Molecular Biology, University of Southern Denmark, Denmark

Lev Levitsky

INEPCP RAS

Moscow, Russia

Elizaveta Solovyeva

INEPCP RAS

Moscow, Russia

Marc Vaudel, University of Bergen

Norway

Stefan Schulze, University of Pennsylvania

USA

Wilhelm, Mathias, Technical University of Munich

Germany

Samaras Patroklos, Technical University of Munich

Germany

Enrique Audaín, University Hospital of Schleswig-Holstein, Kiel,

Germany

Melanie Christine Föll, Northeastern University

USA

Pablo Moreno, EMBL-EBI, UK

European Bioinformatics Institute (EMBL-EBI),

Cambridge, United Kingdom

Johannes Griss, Medical University of Vienna,

Austria

Timo Sachsenberg, Tübingen University

Germany

Mikhail V. Gorshkov

INEPCP RAS

Moscow, Russia

David Bouyssié, IPBS, University of Toulouse, CNRS, UPS

Toulouse, France

Tim Van Den Bossche, Ghent University

Ghent, Belgium

Henry Webel, Novo Nordisk Foundation Center for Protein Research, University of Copenhagen

Denmark

Julian Uszkoreit, Ruhr University Bochum

Bochum, Germany

Eric W. Deutsch

Institute for Systems Biology, USA

Nuno Bandeira

University of California San Diego, USA

Yasset Perez-Riverol

European Bioinformatics Institute (EMBL-EBI)

Cambridge, United Kingdom

**Contributors**

A full list of contributors can be found here: <https://github.com/bigbio/proteomics-metadata-standard#core-contributors-and-collaborators>

**Intellectual Property Statement**

The PSI takes no position regarding the validity or scope of any intellectual property or other rights that might be claimed to pertain to the implementation or use of the technology described in this document or the extent to which any license under such rights might or might not be available; neither does it represent that it has made any effort to identify any such rights. Copies of claims of rights made available for publication and any assurances of licenses to be made available or the result of an attempt made to obtain a general license or permission for the use of such proprietary rights by implementers or users of this specification can be obtained from the PSI Chair.

The PSI invites any interested party to bring to its attention any copyrights, patents or patent applications, or other proprietary rights which may cover technology that may be required to practice this recommendation. Please address the information to the PSI Chair (see contact information at the PSI website).

**Copyright Notice**

Copyright (C) Proteomics Standards Initiative (2021). All Rights Reserved.

This document and translations of it may be copied and furnished to others, and derivative works that comment on or otherwise explain it or assist in its implementation may be prepared, copied, published, and distributed, in whole or in part, without the restriction of any kind, provided that the above copyright notice and this paragraph are included on all such copies and derivative works. However, this document itself may not be modified in any way, such as by removing the copyright notice or references to the PSI or other organizations, except as needed for developing Proteomics Recommendations in which case the procedures for copyrights defined in the PSI Document process must be followed, or as required to translate it into languages other than English.

The limited permissions granted above are perpetual and will not be revoked by the PSI or its successors or assigns.

This document and the information contained herein is provided on an "AS IS" basis and THE PROTEOMICS STANDARDS INITIATIVE DISCLAIMS ALL WARRANTIES, EXPRESS OR IMPLIED, INCLUDING BUT NOT LIMITED TO ANY WARRANTY THAT THE USE OF THE INFORMATION HEREIN WILL NOT INFRINGE ANY RIGHTS OR ANY IMPLIED WARRANTIES OF MERCHANTABILITY OR FITNESS FOR A PARTICULAR PURPOSE."

**Glossary**

All non-standard terms are already defined in detail in section 3.
